# Temperature, but not plant cultivar, influence the efficacy of insecticidal dsRNA in Colorado potato beetle

**DOI:** 10.1101/2024.11.25.625268

**Authors:** Mike Darrington, Jason Solocinski, Sophia K. Zhou, Melise C. Lecheta, Subba Reddy Palli, Yolanda H. Chen, Nicholas M. Teets

## Abstract

Environmental RNAi (eRNAi) is a recent innovation in insect pest control, and comprehensive risk assessment is needed to ensure the environmental safety and longevity of this technology. As eRNAi relies on the insect’s cellular machinery for its mode of action, environmentally mediated plasticity in the activity of cellular processes required for RNAi could influence efficacy and the development of resistance. Here, we investigated the extent to which plant cultivar and temperature influence the efficacy of insecticidal double-stranded RNA (dsRNA) targeting actin in larvae of the Colorado potato beetle (CPB; *Leptinotarsa decemlineata*). Potato cultivar did not significantly affect survival or gene silencing in dsRNA-treated larvae, indicating that efficacy is consistent across potato varieties, at least under laboratory conditions. However, the amount of leaf tissue consumed by larvae varied across cultivars, with consumption being inversely proportional to protein content. Thus, the ingested dose of dsRNA may vary across crop varieties. Temperature did influence RNAi efficacy, with both gene silencing and mortality being reduced when dsRNA treatment occurred at lower temperatures. After three days of feeding with dsRNA, gene silencing occurred at all temperatures, but knockdown efficiency was 62% at 30°C and 35% at 18°C. Beetles consumed significantly less leaf tissue (and thus dsRNA) at 18°C, but the observed temperature-dependent effects could not be fully explained by the quantity of dsRNA ingested. Further, efficacy at different temperatures was not related to transcript levels of core RNAi genes, indicating that other mechanisms are responsible for the observed effects. Overall, these results indicate that environmental conditions can influence the efficacy of insecticidal eRNAi and may affect the rate at which insects develop resistance to these technologies.

## Introduction

RNA interference (RNAi) is a promising method of insect pest control (Baum & Roberts, 2014; Darrington et al., 2017; Palli, 2023). RNAi appropriates a conserved eukaryotic immune response to viral double-stranded RNA (dsRNA), to target specific mRNAs for degradation (Fire et al., 1998). The RNAi pathway is induced when cytoplasmic dsRNAs are processed by Dicer endonucleases into 21bp siRNAs (Hammond et al., 2000; Kim et al., 2009). An Argonaut endonuclease (Carmell et al., 2002) then degrades a single ‘passenger strand’ of duplexed siRNA and binds the remaining ‘guide strand’ to form an RNA-induced silencing complex (RISC) (Tomari et al., 2007). In the final step of RNAi, cellular mRNAs with nucleotide sequences complementary to the RISC’s integral 21bp guide are degraded by the Argonaut’s PIWI domain (Elbashir et al., 2001; Liu et al., 2004). RNAi is an effective control measure when genes that are essential for insects’ performance and survival are targeted for knockdown (Baum & Roberts, 2014; Darrington et al., 2017). Genes such as inhibitor of apoptosis, which regulates cell death (Orme & Meier, 2009), and actin, which is associated with cytoskeletal dynamics (Wolfrum, 1997) and muscle contraction (van Straaten et al., 1999), are good insecticidal targets, but the optimal target for each pest species varies (Cedden et al., 2024; Máximo et al., 2020).

Spraying is the most straightforward system for delivery of insecticidal dsRNAs at scale, although it faces inherent limitations related to dosage and RNA degradation (Christiaens et al., 2020). Spraying also requires that insects initiate RNAi in response to environmentally encountered dsRNA (eRNAi) (Whangbo & Hunter, 2008). Several insects are recalcitrant to eRNAi, but *Coleoptera* are generally susceptible to this technique (Baum & Roberts, 2014; Darrington et al., 2017). Hence, the first sprayable eRNAi technology targeting the Colorado Potato Beetle (CPB; *Leptinotarsa decemlineata*) is now approved for use in the United States (Rodrigues et al., 2021), and advances in the field are anticipated (Palli, 2023). As with all methods of insect control, eRNAi technologies require stringent risk assessments to safeguard public health and manage resistance (Dalakouras et al., 2024; Rodrigues & Petrick, 2020).

As with other insecticides, it appears resistance to eRNAi can evolve quickly. Strains of CPB, western corn rootworm (WCR; *Diabrotica virgifera*) and willow leaf beetle (WLB; *Plagiodera versicolora*) that are highly resistant to eRNAi have been selected for resistance in the laboratory through intergenerational exposure to dsRNA (Khajuria et al., 2018; Liao et al., 2024; Mishra et al., 2021). These reported cases of eRNAi resistance are all caused by autosomal and recessive polymorphisms, but CPB’s resistance is polygenic (Mishra et al., 2021) while WCR’s and WLB’s resistance are monogenic (Khajuria et al., 2018; Liao et al., 2024). The potent (11K-fold) eRNAi resistance observed in CPB (Mishra et al., 2021) may result from selection on a background of pre-existing resistance alleles. This hypothesis is supported by data demonstrating that eRNAi susceptibility varied 7.5-fold for CPB sampled from 14 different sites in Europe (Mehlhorn et al., 2020). Furthermore, CPB’s resistance to chemical insecticides developed largely through selection on standing variation (Pélissié et al., 2022).

Resistance or reduced susceptibility to eRNAi can be attributed to numerous underlying mechanisms (Darrington et al., 2017). Failure of systems that transport and process dsRNA from the environment appear to be a primary factor influencing insects’ sensitivity to eRNAi (Yoon et al., 2016). Several insect taxa that are susceptible to eRNAi transport dsRNA within endosomes (Cappelle et al., 2016; Xiao et al., 2015; Yoon et al., 2016), which mature and release their contents following a process of V-ATPase induced acidification (Lafourcade et al., 2008). For some insects, VHA16, a subunit of the V-ATPase transmembrane domain (Vₒ), triggers the release of dsRNA from endosomes and is essential for eRNAi (Cappelle et al., 2016; Saleh et al., 2006; Yoon et al., 2016). For CPB, eRNAi also depends on StaufenC, an RNA-binding protein that enhances Dicer’s activity (Kim et al., 2021) and facilitates transport of dsRNA through the endoplasmic reticulum (Koo & Palli, 2024). Indeed, both WCR and WLB developed eRNAi resistance when systems of dsRNA transport lost function (Khajuria et al., 2018; Liao et al., 2024). The activity and localization of nucleases that digest dsRNA can also influence susceptibility. Insects from several orders, including aphids (Christiaens et al., 2014), locusts (Song et al., 2017) and moths (Wang et al., 2016) are recalcitrant to eRNAi as a result of nucleases that degrade effector dsRNA.

The above processes that influence susceptibility to eRNAi could also be influenced by environmental conditions. However, the extent to which the environment modulates the efficacy and evolution of resistance to insecticides, particularly eRNAi, has received little attention. Thus, understanding how eRNAi functions in variable environments is essential for optimizing its efficacy and minimising the risk of resistance. Environmentally mediated reductions in eRNAi efficacy could accelerate the evolution of resistance by increasing the population size available for *de novo* mutations and facilitating an accumulation of alleles involved in quantitative resistance. Further, evolutionary processes such as genetic assimilation provide a route for environmentally plastic phenotypes to be canalized in a population (Ghalambor et al., 2007). Indeed, genetic assimilation appears to have played a role in evolved tolerance to insecticide exposure in wood frog populations living in agricultural ecosystems (Hua et al., 2015).

The quality of an insect’s diet can moderate the pathogenicity of viral infections (Ballard et al., 2000; Lasa et al., 2009; Lee et al., 2006), suggesting a link between nutrition and the activity of RNAi. However, the precise effects of nutrition appear to vary across species. Beet armyworm (*Spodoptera exigua*) is more susceptible to viral infection when fed a high protein diet (Lasa et al., 2009), while the congeneric African cotton leafworm (*Spodoptera littoralis*) mounted a robust anti-viral response when given supplementary protein (Lee et al., 2006). In the coddling moth (*Cydia pomonella*) viral immunity was primarily related to carbohydrate intake rather than protein (Ballard et al., 2000). At the molecular level, the forkhead box O (FOXO) family of transcription factors, which are activated in times of nutritional stress (Zhang et al., 2018), may provide the link between nutrition and RNAi. FOXO represses viral infections in silk moth (Kang et al., 2021) and regulates the expression of both Dicer and Argonaut in *Drosophila melanogaster* (Spellberg & Marr, 2015). If insects’ eRNAi responses are in fact nutrition dependent, the variable nutrient profiles across crop cultivars (Domek et al., 1995; Khorramifar et al., 2023; Lee et al., 2016; Minhas et al., 2004) may have implications for eRNAi based management for plant feeding pests.

Insects’ responses to eRNAi may also be modulated by variations in environmental temperature. Since temperature directly affects insects’ metabolism and enzyme kinetics (Gillooly et al., 2001), pest species may may consume and process more dsRNA from the environment at elevated temperatures. Moreover, heat-shock proteins, which can be activated at elevated temperatures (Chen et al., 2020), have been shown to inhibit the RNAi pathway in the diamondback moth *(Plutella xylostella*) (Lin et al., 2023). In the context of viral immunity, lower temperatures reduce the activity of RNAi and make mosquitoes susceptible to infection (Adelman et al., 2013). Similarly, in both flatworms exposed to eRNAi and fruit flies that transgenically express dsRNA, RNAi efficiency is reduced at lower temperatures (Wudarski et al., 2019; Zhang et al., 2020). Temperature could also influence the evolution of resistance to dsRNA, as resistance-related mutations to imidacloprid arise on a temperature-dependent basis in *D. melanogaster* (Fournier-Level et al., 2019). However, the extent to which temperature influences the efficacy of insecticidal dsRNA has not been assessed.

Here, we investigated the extent to which plant cultivar and temperature affect responses to environmental dsRNA in CPB, to identify environmental factors that might reduce efficacy and contribute to resistance development. CPB is a well-established pest that is resistant to at least 50 chemical insecticides (Alyokhin et al., 2008), is the target of a commercially available eRNAi product (Rodrigues et al., 2021), and has previously developed eRNAi resistance under stable laboratory conditions (Mishra et al., 2021). We assessed the efficacy of insecticidal eRNAi in different environments by quantifying knockdown efficiency of the target gene and survival. Larvae were fed dsRNA targeting actin on either three different potato cultivars at 25°C, or at 18, 25 or 30°C on a common potato cultivar. To determine if gene silencing and survival across potato cultivars was related to macronutrient profile, we quantified protein, carbohydrate, and lipid content for each potato cultivar. We also quantified consumption of leaf tissue on each variety and at each temperature, to determine whether the efficacy of eRNAi in different treatments was explained by rates of consumption. Finally, we compared the expression of key eRNAi genes across temperatures, to elucidate potential mechanisms that might explain that variable efficacy.

## Results

### Effect of potato cultivar on survival

Treatment with dsRNA targeting actin (dsactin) negatively impacted survival (ANOVA, df=1, *χ*^2^=58.2, p<0.001), but there was no significant interaction between dsactin treatment and cultivar (ANOVA, df=2, *χ*^2^=2.6, p>0.05) (Fig 1), indicating that responses to eRNAi were similar across plant cultivars. In the untreated controls, survival improved by 13% when larvae were fed Chieftain potatoes and Purple Viking potatoes, in comparison to Kennebec potato, although these effects were not significant (ANOVA, df=2, *χ*^2^=0.42, p>0.05).

**Figure 1.**
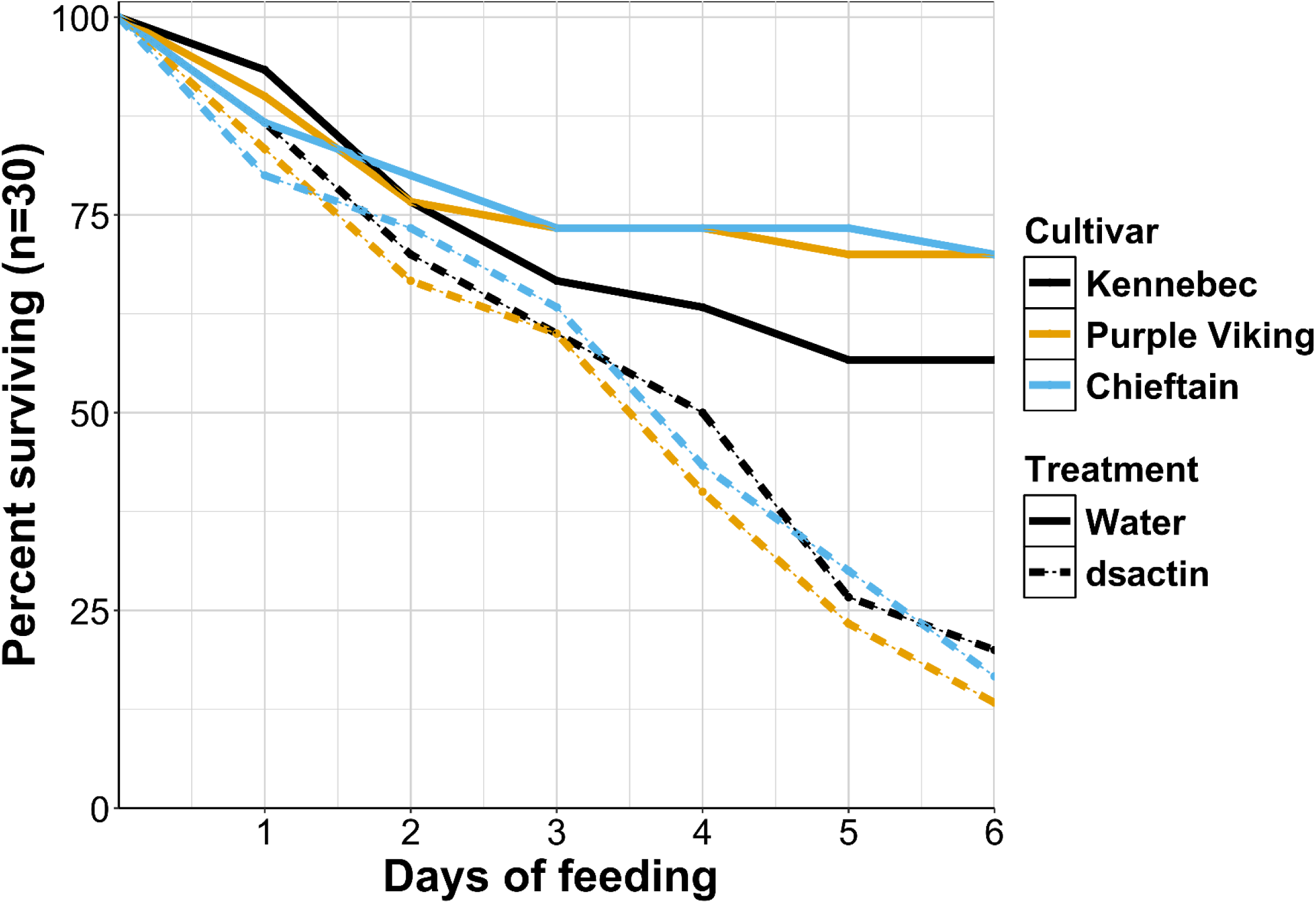
Survival of CPB larvae on three potato cultivars. Larvae were treated with either dsactin, or water for three days. After six days of observation, survival was significantly lower for beetles fed dsRNA (ANOVA, df=1, χ^2^=58.2, p<0.001), but survival was not significantly affected by cultivar (ANOVA, df=2, χ^2^=0.42, p>0.05).

### Effect of potato cultivar on gene silencing over time

In untreated controls, actin mRNA abundance increased slightly between the 24- and 72-hour treatment groups (Fig 2). Treatment with dsactin significantly reduced actin expression (ANOVA, df=1, F=25.2, p<0.001), but the gene silencing effect of eRNAi was not significantly different between cultivars, as indicated by a nonsignificant interaction between dsactin treatment and cultivar (ANOVA, df=2, F=0.13, p>0.05). Actin expression was significantly affected by an interaction between cultivar and feeding period (ANOVA, df=4, F=3.81, p<0.01), indicating some cultivar-specific variation in actin mRNA abundance over time. For larvae fed Kennebec potato, dsactin treatment reduced actin mRNA abundance by 16.7% after 24 h, 54.3% after 48 h and 41.5% after 72 h, relative to untreated controls. When larvae were fed Chieftain potato, dsactin treatment reduced actin mRNA abundance by 35.4% after 24 h, 32.6% after 48 h and 62.3% after 72 h. For larvae fed Purple Viking potato, actin mRNA abundance was reduced by 18.8% after 24 h, 41.1% after 48 hours and 57.1% after 72 hours, when larvae were fed dsactin.

**Figure 2.**
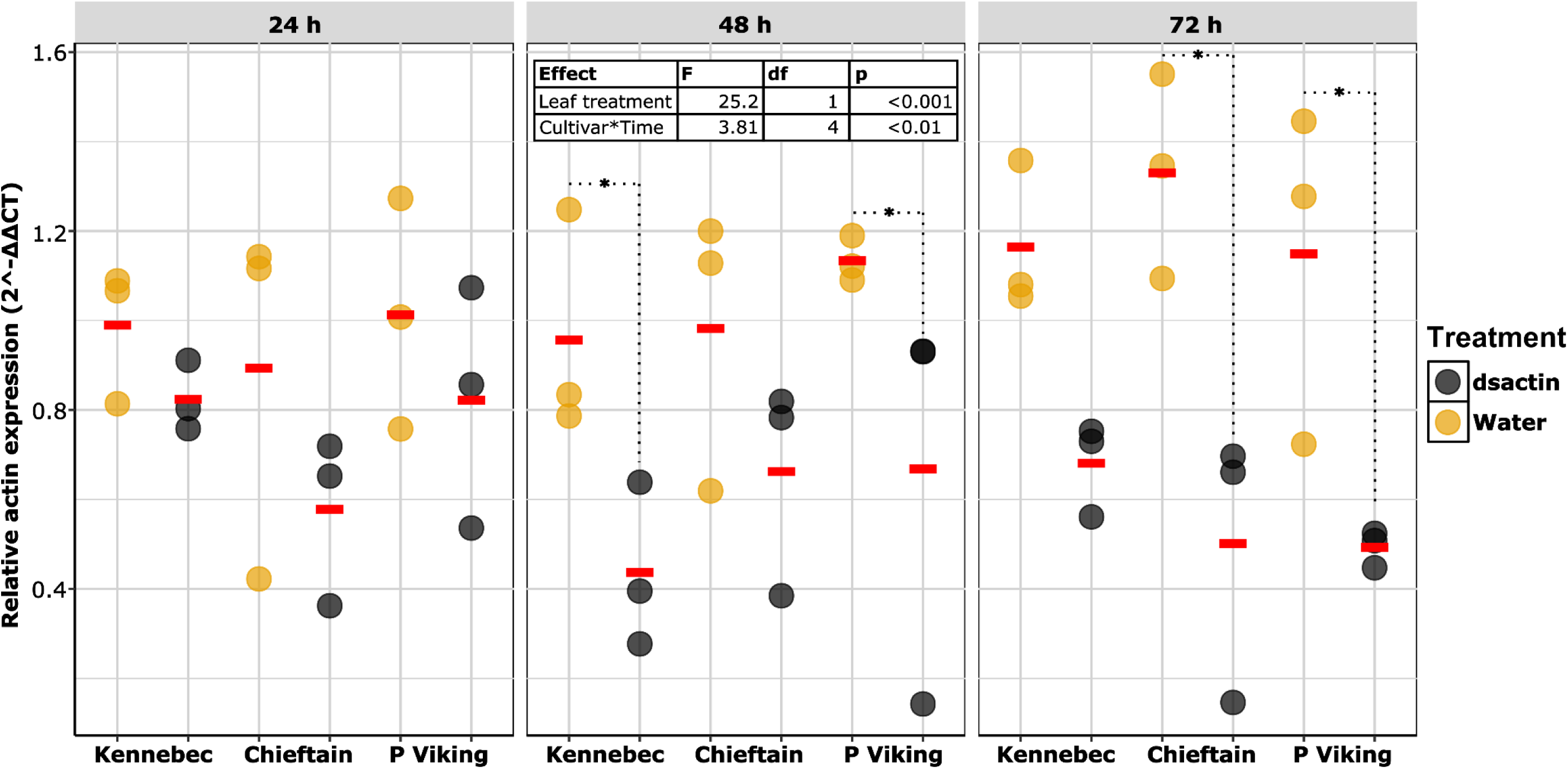
Actin expression of CPB larvae fed three different potato cultivars. Larvae were treated with either dsactin or water for three days. Mean actin expression for each group is represented by a red bar. dsactin treatment had a significant effect on gene silencing. Cultivar and feeding period interacted to have a significant effect on gene silencing. Groups of larvae with significant knockdown at a particular combination of time point and cultivar are indicated by an asterisk (emmeans, sidak, p<0.05). The table at the top summarizes the factors that had a significant effect on actin transcript abundance.

### Between cultivar variation in metabolite content

Colorimetric analyses were carried out to compare the macronutrient content of leaves from Kennebec, Chieftain and Purple Viking potatoes. Potatoes exhibited significant between-cultivar differences in protein (ANOVA, df=2, F=10.5, p<0.001), soluble carbohydrates (ANOVA, df=2, F=6.7, p<0.001) and starch content (ANOVA, df=2, F=15.2, p<0.0001), with Purple Viking being the most nutrient dense cultivar (Table 1). Purple Viking contained significantly more soluble carbohydrates (ANOVA, Tukey, p<0.05) and starch (ANOVA, Tukey, p<0.01) than both Kennebec and Chieftain, and significantly more protein than the Kennebec strain (ANOVA, Tukey, p<0.0001). There were no significant differences in lipid content between cultivars (ANOVA, df=2, F=1.14, p>0.05).

**Table 1.** Metabolite content of leaf tissue from three distinct potato cultivars. Values indicate the mean nutrient content of 113 mm^2^ leaf discs from each cultivar (n=10). Superscript letters indicate significant differences between cultivars (ANOVA, Tukey, p<0.05).

| Cultivar | Mean protein (µg) | Mean soluble carbs (µg) | Mean starch (µg) | Mean lipids (µg) |
| --- | --- | --- | --- | --- |
| Kennebec | 1.42 (SE 0.13)<sup>a</sup> | 2.56 (SE 0.45)<sup>a</sup> | 20.7 (SE 0.56)<sup>a</sup> | 57.2 (SE 7.06)<sup>a</sup> |
| Chieftain | 1.81 (SE 0.08)<sup>ab</sup> | 2.12 (SE 0.56)<sup>a</sup> | 17.4 (SE 0.69)<sup>a</sup> | 48.3 (SE 4.55)<sup>a</sup> |
| Purple Viking | 2.33 (SE 0.18)<sup>b</sup> | 5.13 (SE 0.81)<sup>b</sup> | 25.2 (SE 1.47)<sup>b</sup> | 63.4 (SE 8.38)<sup>a</sup> |

### Area of leaf consumed by larvae on different potato cultivars

L1 larvae were contained for 24 hours in 12 mm diameter (ø) clip cages on the leaves of Kennebec, Chieftain and Purple Viking potatoes that had been treated with dsactin or water. There were no significant differences in leaf consumption between larvae fed dsactin or water on a common cultivar, suggesting that dsRNA itself does not have an immediate antifeedant effect for CPB larvae. There were significant between-cultivar differences in leaf consumption (ANOVA, df=2, F=6.3, p<0.001), primarily between the Kennebec and Purple Viking cultivars (ANOVA, Tukey, p<0.001). The area of leaf tissue consumed in 24 hours ranged from 13.6 – 52.5 mm^2^ for the Kennebec cultivar, 12.7 – 45.2 mm^2^ for the Chieftain cultivar and 2.9 – 39.9 mm^2^ for the Purple Viking cultivar (Fig 3).

**Figure 3.**
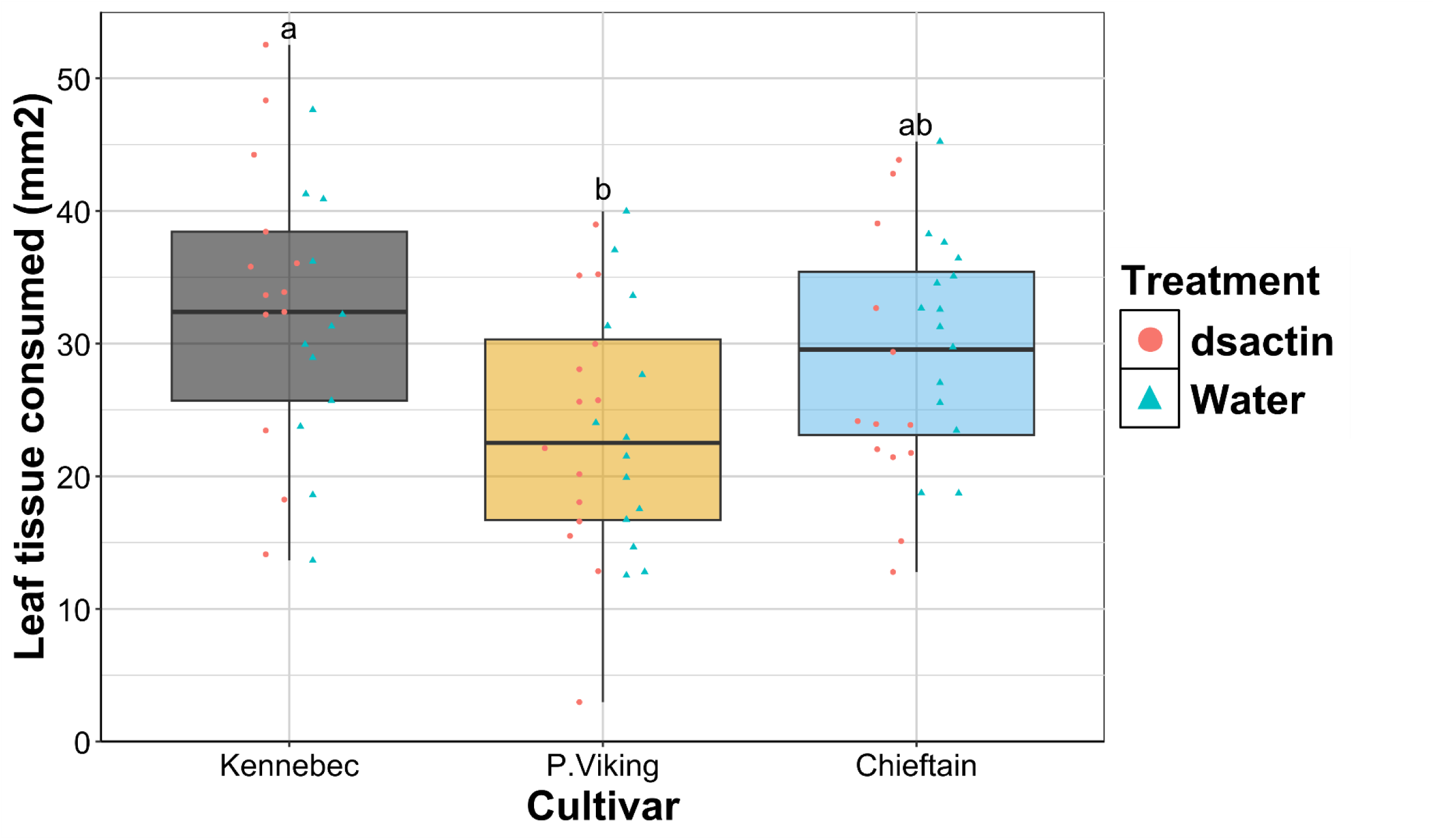
Area of leaf tissue consumed by first instar larvae in 12 mm *ø* clip cages on Kennebec, Chieftain or Purple Viking potato. Either water or dsRNA was applied to leaf surfaces before larvae were trapped on those leaves for 24 hours. Consumption did not differ significantly when leaves were treated with dsRNA but there were significant between-cultivar differences in consumption (ANOVA, df=2, F=6.3, p<0.001). Thus, the dsactin and water-treated leaves are plotted together for each cultivar. Letters denote significant differences between groups revealed by post-hoc analysis (Tukey HSD, p<0.05).

### Effect of temperature and dsactin treatment on survival

First instar larvae were kept on leaves coated with either dsactin or water for three days at 18, 25, or 30°C, after which larvae were kept at these temperatures on untreated leaves, and mortality was recorded until day six (Fig 4). At 18°C, 36% (SE 8.8) of larvae fed dsactin and 80% (SE 5.8) of larvae fed water survived. At 25°C, 3% (SE 3.3) of larvae survived dsactin treatment and 66% (SE 8.8) of larvae survived water treatment. All larvae fed dsactin at 30°C had died after four days, while 43% (SE 8.8) of larvae fed water survived the 6-day observation period. Both temperature (ANOVA, df=2, *χ*^2^=77.9, p<0.001) and dsactin treatment (ANOVA, df=1, *χ*^2^=77.4, p<0.001) had significant effects on survival, and there was also a significant interaction between dsactin treatment and temperature that affected survival (ANOVA, df=2, *χ*^2^=6.23, p<0.05).

**Figure 4.**
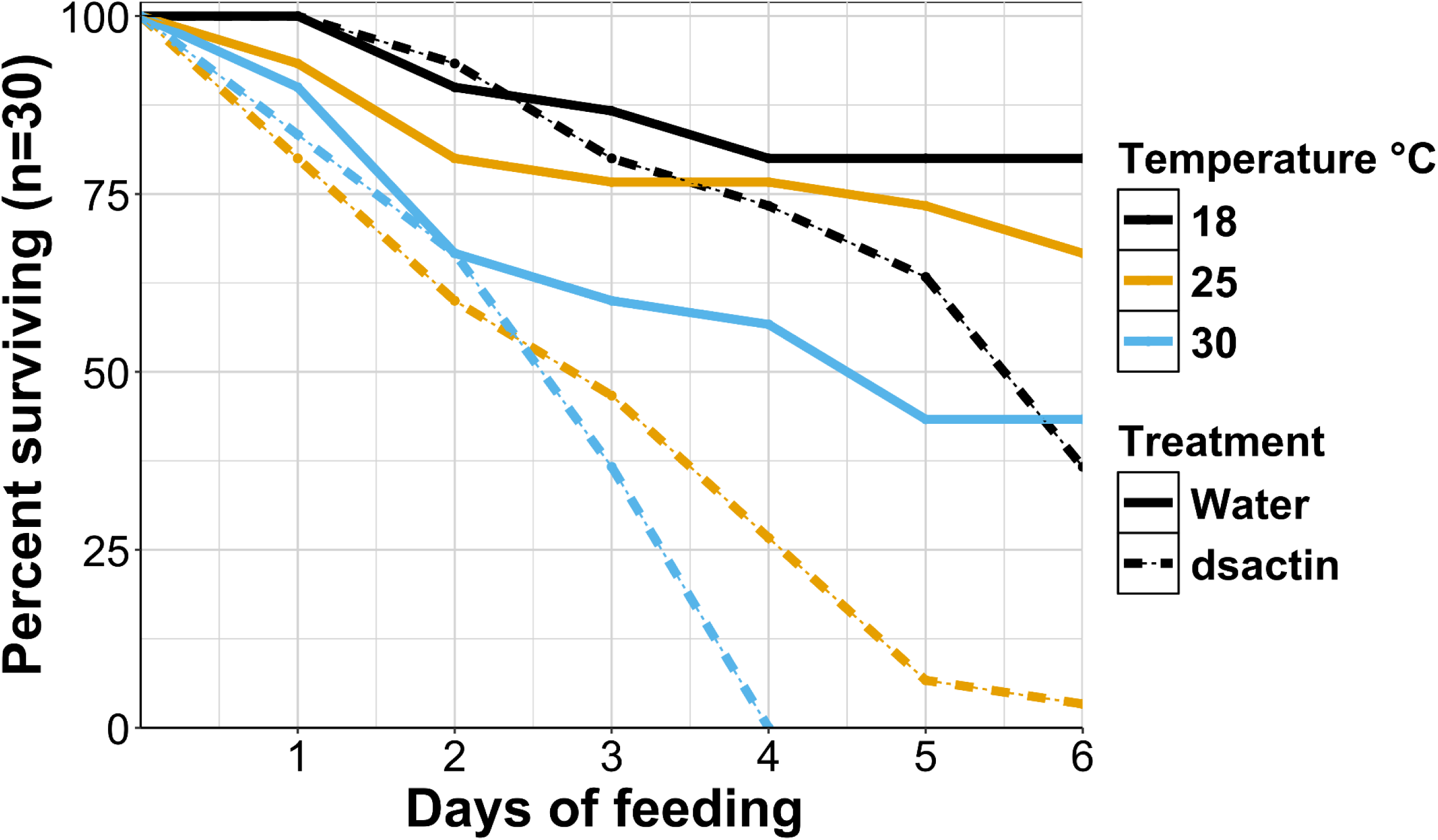
Survival of CPB larvae treated with actin dsactin at three temperatures. Larvae were treated with either dsactin or water for three days at 18, 25, or 30°C, and observed for an additional three days. Survival was significantly lower when beetles were exposed to higher temperatures (ANOVA, df=2, χ^2^=77.9, p<0.001) or fed dsRNA (ANOVA, df=1, χ^2^=77.4, p<0.001). Temperature and feeding period interacted significantly to affect survival outcomes (ANOVA, df=2, χ^2^=6.23, p<0.05).

### Effects of temperature, dsactin and feeding period on gene silencing

First instar larvae were fed leaves coated with either dsactin or water for three days at 18, 25 or 30°C. The relative expression of actin was then measured by RT-qPCR for each group after 24, 48 and 72 hours (Figure 5). After 72 hours, actin mRNA titers were comparable at all temperatures for larvae fed on dsactin but varied for larvae fed on water (Fig 5). Knockdown efficiency increased with temperature and over time, with the effects of temperature (ANOVA, df=2, F=18.2, p<0.001), treatment with dsactin (ANOVA, df=1, F=57.6, p<0.001) and feeding period (ANOVA, df=2, F=23.7, p<0.001) all being significant. Temperature and feeding period also interacted significantly to affect actin expression (ANOVA, df=4, F=29.2, p<0.001). When larvae were fed dsactin at 18°C, actin mRNA abundance was reduced by 12.9% after 24 h, 35.5% after 48 h and 35.6% after 72 h. At 25°C, dsactin treatment reduced actin mRNA by 23.8% after 24 h, 55.3% after 48 h and 45.4% after 72 h. For larvae reared at 30°C, actin mRNA abundance was reduced by 42.5% after 24 h, 32.3% after 48 hours and 62.4% after 72 hours, when larvae were fed dsactin.

**Figure 5.**
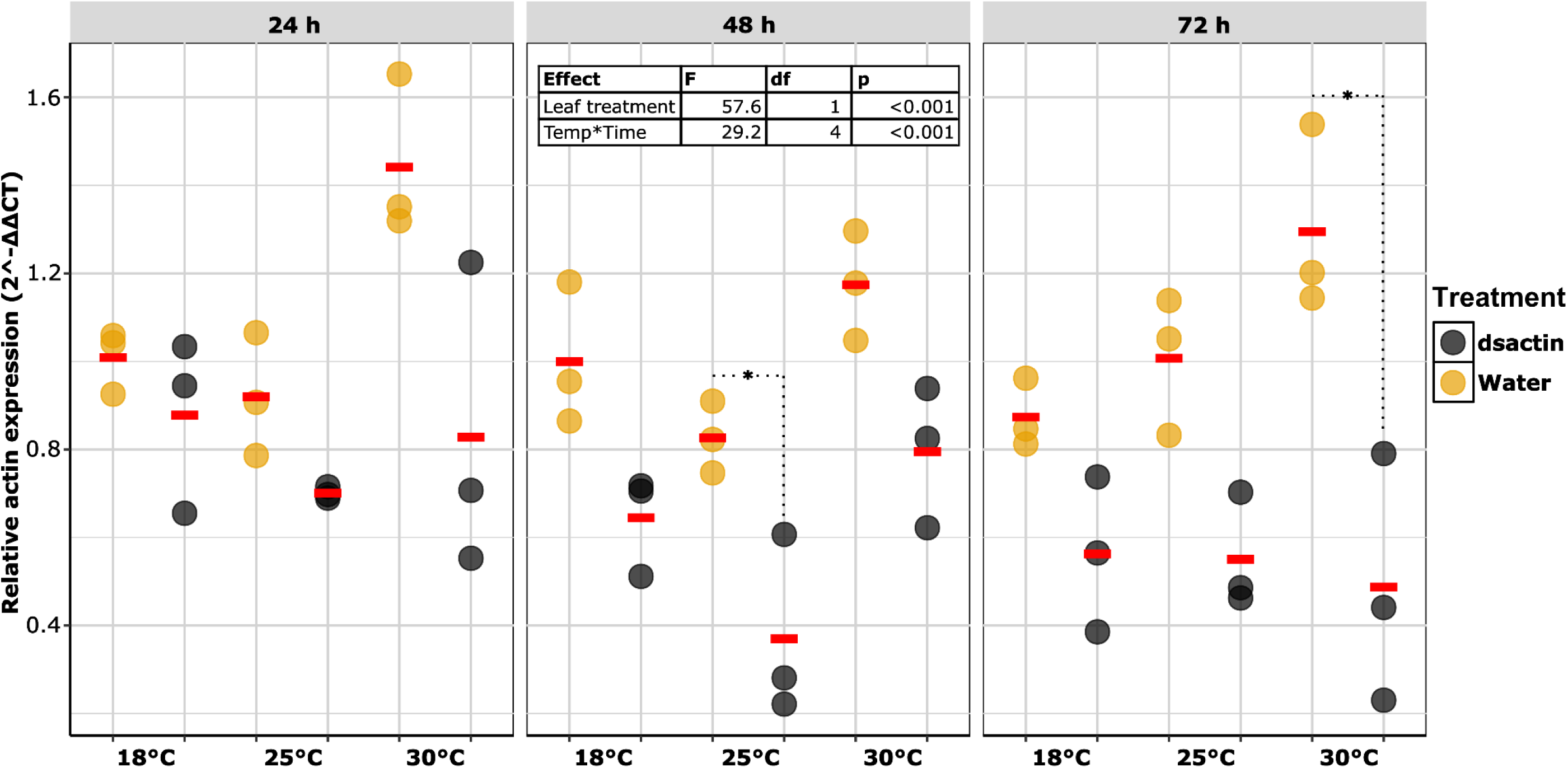
Relative actin expression for CPB larvae held at either 18, 25 or 30°C while fed either dsactin or water for 72 hours. Mean actin expression for each group is represented by a red bar. dsactin treatment had a significant effect on gene silencing. Temperature and feeding period interacted to have a significant effect on gene silencing. Groups of larvae with significant knockdown at a particular combination of time point and temperature are indicated by an asterisk (emmeans, sidak, p<0.05).

### Effect of temperature on potato consumption

Treatment with dsactin did not significantly affect larval consumption of leaf tissue at 18, 25 or 30°C. There were significant temperature dependent differences in leaf consumption (ANOVA, df=2, F=40.7, p<0.0001). After 24 hours at 18°C, larvae had consumed between 6.1 – 27.6 mm^2^ of leaf tissue, which was significantly less than the amount of tissue consumed at 25° and 30°C (ANOVA, Tukey, p<0.0001). The area of leaf tissue consumed was not significantly different between the 25°C (18.9 – 60 mm^2^) and the 30°C treatments (20.8 - 86.3 mm^2^) (Fig 6).

**Figure 6.**
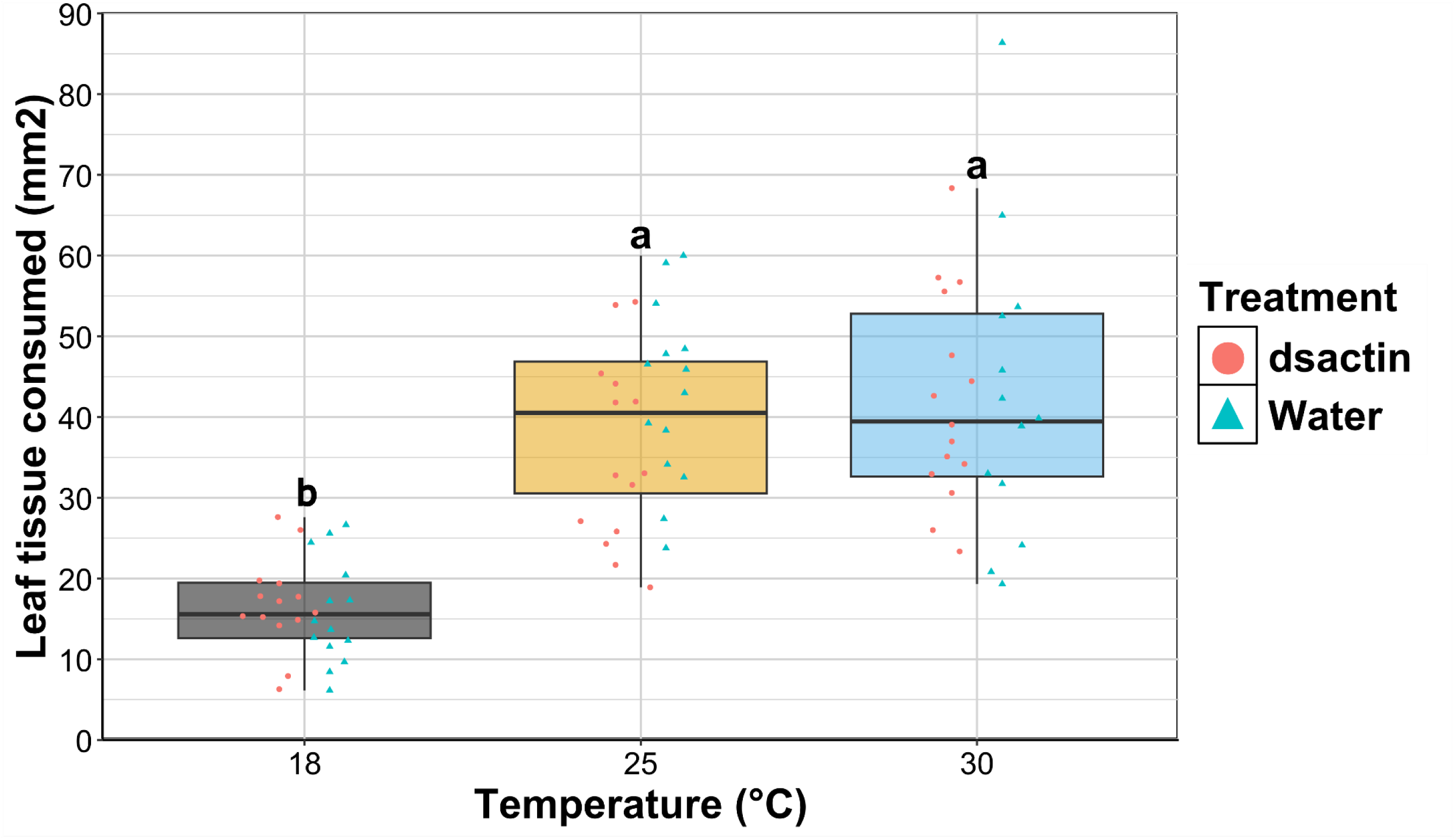
Leaf tissue consumed by CPB larvae that were held at either 18, 25 or 30°C while being treated with either dsactin or water for 24 hours. dsactin treatment did not have a significant effect on leaf consumption at any temperature (lm, df=5,78, t=0.11, p>0.5). Thus, both groups are plotted together for each temperature. Larvae consumed significantly less leaf tissue at 18°C than at 25° or 30°C (ANOVA, Tukey, p<0.0001).

### Effect of temperature on the expression of eRNAi associated genes

Treatment with dsactin for 24 hours did not significantly affect the expression of core RNAi genes at any temperature (Figure 7). Relative to treatment at 25°C, Dicer mRNA abundance was significantly reduced at 18°C (lm, t=2.7, p<0.05) and 30°C (lm, t=2.5, p<0.05). Treatment at 18°C (lm, t=2.3, p<0.05) but not 30°C significantly reduced the abundance of Argonaut mRNA. Relative to 25°C, StaufenC mRNA abundance was significantly elevated at 18°C (lm, t=-2.4, p<0.05) but not 30°C. VHA16 mRNA abundance did not differ significantly at any temperature.

**Figure 7.**
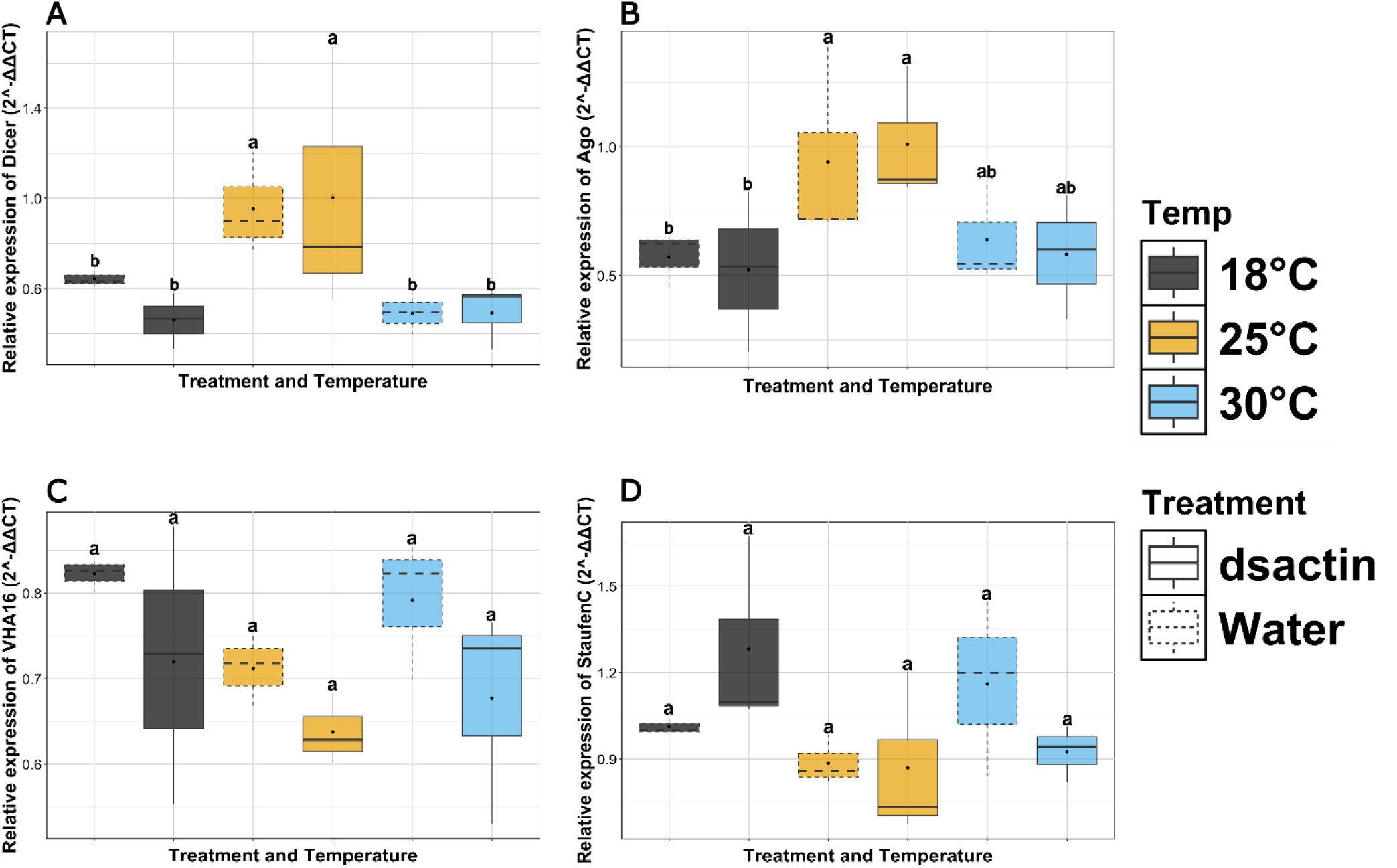
Relative mRNA abundance of core eRNAi genes in response to dsactin treatment at different temperatures. Expression of Dicer (A), Argonaut (B), VHA16 (C) and StaufenC (D) was measured in CPB larvae fed dsRNA or water while being held at either 18°C, 25°C or 30°C for 24 hours. After 24 hours, treatment with dsactin did not significantly affect the expression of any gene, at any temperature. Temperature significantly affected relative expression of Dicer, Argonaut and StaufenC. Letters denote significant differences in revealed by Post-hoc analysis (Tukey HSD, p<0.05).

## Discussion

Insect control products using eRNAi are now commercially available (Head et al., 2014; Rodrigues et al., 2021), and it has been demonstrated that insects can develop resistance in the laboratory (Khajuria et al., 2018; Liao et al., 2024; Mishra et al., 2021). While there is a clear genetic component in determining susceptibility to eRNAi (both within and across species; see (Darrington et al., 2017)), the extent to which environmental conditions influence the efficacy of this technology has not been assessed. Understanding the contribution of the environment to eRNAi responses is critical for maximizing control and managing resistance. Here, we investigated the extent to which diet and temperature affect eRNAi for CPB, as conditions that reduce the toxicity of insecticidal dsRNA may permit the maintenance of larger pest populations and increase the risk of resistance.

While the consumption of dsRNA-treated leaves varied across potato cultivars, rates of mortality were consistent. In our experiments, the Purple Viking cultivar contained significantly more soluble carbohydrates and starch than both the Kennebec and Chieftain cultivars. Furthermore, Purple Viking contained significantly more protein than Kennebec. Differences in macronutrient content corresponded with differences in leaf consumption, with consumption generally being higher in leaves with lower macronutrient density. This result is consistent with the nutritional geometry framework, which posits that herbivores continue foraging until reaching an intake target for a limiting nutrient, typically protein (Raubenheimer & Simpson, 2018), and it indicates that the realized dose of dsRNA in the field may depend on plant nutrient profile. As mortality did not vary across cultivars in our experiments, despite differences in consumption, this suggests that either dsactin is sufficiently toxic to compensate for the variation in dose, or that the eRNAi response is intensified under high nutrition conditions. While we cannot tease apart these possibilities in our experiments, previous work has shown a link between nutrition and antiviral immunity (Ballard et al., 2000; Lasa et al., 2009; Lee et al., 2006). For example, multiply-enveloped nucleopolyhedrovirus (SeMNPV) was significantly more lethal when beet armyworm (*Spodoptera exigua*) larvae had been fed supplementary protein (Lasa et al., 2009). In contrast, when fed supplementary protein, the African cotton leafworm (*Spodoptera littoralis*) exhibited enhanced antiviral function via improved lysozyme activity and encapsulation responses (Lee et al., 2006). In a field setting, polyphagous insects like CPB can consume a range of host plants with varying macronutrient profiles, which could in turn affect the rate at which they consume dsRNA-treated plants and induce RNAi. Future studies using artificial diets with specified macronutrient profiles could directly test the links between nutrients, rates of consumption, and activity of the RNAi pathway.

While RNAi efficacy was consistent across plant cultivars, temperature had significant effects on both survival and gene silencing. Indeed, while nearly 40% of dsactin-treated larvae survived the six-day observation period at 18°C, survival was at or near 0% in the groups treated at 25 and 30°C. The reduced survival at 18°C was reflected by lower knockdown efficiency, as knockdown efficiency 72 h after treatment began was far lower at 18°C (~35%) than at 30°C (~63%). One limitation of our study is that leaf disc assays were restricted to a six-day period of observation, as at that point larvae had reached the L3 stage and became too large for the bio-assay trays. Survival in the 18°C treatment was trending downward at the end of the study, and thus mortality may have continued to increase if beetles were allowed to develop further. Nonetheless, lower temperatures clearly delayed mortality, and beetles fed dsactin at 18°C continued to feed and grow for the duration of the experiment (data not shown), suggesting increased plant damage could occur at lower temperatures prior to the onset of mortality. While we were unable to measure consumption in our leaf disc assays, due to discs becoming distorted by the end of the observation period, in ongoing experiments we are assessing these responses on whole plants to see if beetles can complete life cycle and accurately estimate total herbivory.

The most likely explanation for reduced eRNAi efficacy at 18°C is reduced ingestion of dsRNA. Beetles at 18°C consumed less than half the amount of leaf tissue over the first 24 h than those at 25 and 30°C. While this result is not surprising, due to reduced metabolic demands at lower temperatures, it does have implications for field application of dsRNA. Due to washout and instability, dsRNA applications may deteriorate over time at any temperature (Bachman et al., 2020), but if spraying occurs during periods of lower temperatures, beetles may escape lethality while the dsRNA is still present. In addition to reduced consumption, previous studies have demonstrated reduced activity of RNAi at lower temperatures, independent of dosage (Adelman et al., 2013; Wudarski et al., 2019; Zhang et al., 2020). Thus, reduction in dosage at lower temperatures may be compounded by reduced activity of RNAi machinery. Consistent with this idea, while consumption was equivalent at 25 and 30°C, beetles at 30°C died faster and had increased gene knockdown, suggesting higher activity of the RNAi machinery at 30°C.

To test for evidence that environmental conditions directly influence the RNAi pathway itself, we measured transcript abundance of a core panel of RNAi genes described by Yoon et al. (2016). There was a trend towards lower expression of Dicer and Argonaut at 18 and 30°C relative to 25°C, with StaufenC having the opposite pattern, although several of these changes were not statistically significant. Regardless, these results suggest that core eRNAi genes do not have a linear relationship with temperature, at least at the transcript level. One other study has investigated transcriptional regulation of RNAi machinery as a function of temperature in the oriental fruit fly (*Bactrocera dorsalis*), which significantly downregulated Dicer and Argonaut in response to severe heat stress (Xie et al., 2017). While we did not observe significant downregulation of Argonaut at higher temperatures, the temperatures used in our experiments were within the permissible growth range of CPB. However, temperature-dependent regulation of the RNAi pathway warrants further investigation, as Dicer is subjected to complex posttranslational regulation (Kurzynska-Kokorniak et al., 2015). Additional work should also consider whether extreme events affect insecticidal RNAi, given previously identified links between stress proteins and RNAi activity (Lin et al., 2023).

The expression patterns of actin across temperatures may have also contributed to the observed variation in eRNAi efficacy. Actin had elevated transcript abundance at higher temperatures, and RNAi can be more efficient when the expression of target genes is elevated (Chen et al., 2021). Thus, identifying target genes with stable expression across temperatures may lead to more consistent eRNAi outcomes in varied climates. For CPB, dsRNA targeting inhibitor of apoptosis is more effective than actin under standard laboratory conditions (Máximo et al., 2020), and a full factorial experiment systematically manipulating temperature and target gene could be used to identify candidate genes that perform consistently across an environmental gradient.

## Conclusions

Our results indicate that environmental factors should be considered when developing eRNAi products and evaluating their potential ecological risks. The toxicity and knockdown efficiency of dsRNA was consistent across three potato cultivars, suggesting plant variety is unlikely to influence field outcomes or the development of resistance. However, cultivars did have varied macronutrient profiles and rates of consumption, and given previously identified links between nutrition and antiviral activity, care should be taken when designing field application protocols for dsRNA insecticides. Specifically, the potential effects of host plant variety on the ingested dose and activity of RNAi should be considered to ensure equal outcomes across common cultivars.

Consistent with prior research, insecticidal dsRNA was less toxic at low temperatures, suggesting that eRNAi might be more likely to produce resistant phenotypes if applied in a colder climate. Successful eRNAi requires several steps, including ingestion, transport and uptake to target tissues, and processing by the RISC complex, and some or all of these processes could be temperature-dependent. While reduced consumption is the likely explanation for our results, additional work is needed to tease apart the precise mechanism(s) that underpin reduced eRNAi efficacy at low temperatures. Follow-up experiments are also underway to determine whether beetles treated with dsRNA at low temperatures can complete the life cycle and to quantify total herbivory at each temperature, to better simulate potential field outcomes.

## Methods

### Potato horticulture

Three potato cultivars that are commonly grown were selected to represent a range of plant nutrition that beetles will encounter in the field. Organic Kennebec, Chieftain and Purple Viking potato seeds were supplied by The University of Kentucky Community Supported Agriculture Project. Potatoes were grown in unfertilised Miracle Grow potting soil, in separate six-inch pots, in a controlled greenhouse (25°C ±5; 16L:8D).

### Insect husbandry

We maintained a CPB colony originally collected from organic farms in Vermont, USA. Since eRNAi has not been approved for organic production, these beetles have not been previously exposed to eRNAi. Beetles were reared in a controlled greenhouse (25°C ±5; 16L:8D) on organic potato plants (4-6 weeks old). Eggs were collected from the plants as needed and moved to an incubator (25°C; 16L:8D) where they were stored on moist tissue in a Petri dish until hatching.

### eRNAi bioassays

We fed CPB larvae leaves coated with insecticidal dsactin or water as a control and quantified survival and gene silencing outcomes, while manipulating two environmental factors: 1) the potato cultivar treated with insecticidal dsRNA, and 2) ambient temperature. The direct effects of cultivar and temperature on gene silencing and survival were investigated independently, using an equivalent balanced experimental design. Three levels of each factor (cultivar or temperature) were analysed for the two leaf treatments (dsactin or water), giving six feeding groups for each factor. The experiment was replicated three times for each factor, and each replicate consisted of six feeding groups of 16 larvae. To assess the direct effects of cultivar, larvae were fed leaf disks from Kennebec, Chieftain or Purple Viking potatoes at 25°C. To assess temperature effects, larvae were exposed to either 18, 25 or 30°C and given only Kennebec potato. To begin, 10 mm diameter (Ø) leaf discs were placed in independent cells (12 mm Ø X 15 mm height) of a 128 well a Bio-assay tray (Frontier). 4 µl aliquots of either insecticidal dsRNA (1:1, 900 ng/µl dsactin: Triton-X (53 nM)), or water (1:1, H2O: Triton-X (53 nM)) were pipetted onto the discs, and a single first instar larvae was immediately placed in each cell. The trays were then incubated at the appropriate temperature (16L:8D, 69% RH) in programmable incubators. On days two and three, a subset of two larvae were taken from each feeding group for RT-qPCR, and remaining larvae were placed on fresh discs that had been treated with either dsRNA or water as appropriate. On day four, two larvae were extracted from each feeding group for RT-qPCR, and at this point (72 h after the onset of the experiment) the dsRNA treatment concluded. For days 4-6, larvae were kept on untreated leaf discs on the same cultivar or at the same temperature, and survival continued to be monitored. The experiment ran for six days with survival for each feeding group being scored every 24 hours from day one. The experiment was concluded at day six because surviving larvae grew too large for the bioassay trays.

### RNA extraction, cDNA synthesis and RT-qPCR

#### RNA extraction

RNA for both dsRNA synthesis and RT-qPCR was extracted from groups of two larvae. The larvae were homogenised in 400 µl of TRI Reagent (Invitrogen) using a FastPrep-24 bead beater (MP) and two, 2.3 mm zirconia/silica beads (Biospec). Phase separation was carried out by mixing the homogenised solution with chloroform (VWR) (5:1) and centrifuging for 12000 g and 4°C for 15 mins. The aqueous phase was then mixed ≈1:1 with 100% isopropanol (VWR) and incubated at −80°C for two hours. RNA was pelleted by centrifugation at 12000 g for 10 mins and 4°C, and the pellet was washed with 75% Ethanol (VWR). The washed pellet was centrifuged at 7500 g and 4°C for 5 mins, and the wash step was then repeated. RNA was assessed for purity and concentration with a CLARIOstar plate reader (BMG).

#### dsRNA synthesis

A 305bp dsRNA fragment targeting actin was synthesised using minor revisions to the protocol published by Zhu et al. (2011). The target fragment was PCR amplified from cDNA using Taq 2X mastermix (NEB). PCR products were then purified in triplicate, using a single QIAquick PCR purification kit (Qiagen) column. 30 µl preparations of dsRNA were synthesised from 60 ng of purified PCR product with the HiScribe T7 Quick High Yield RNA Synthesis Kit (NEB). The products were treated simultaneously with DNAse1 (NEB) and RNase A (Promega) before being precipitated in 3 M sodium acetate (0.1 V/V; pH 5.2) and 100% ethanol (2.5 V/V) at −80°C. The dsRNA was pelleted by centrifugation at 19000 g and 4°C for 15 mins, washed with 70% ethanol, washed again and air dried. dsRNA purity and concentration were measured using a CLARIOstar plate reader (BMG).

#### RT-qPCR

cDNA was reverse transcribed from 50 ng of RNA per replicate, with a RevertAid first strand synthesis kit (ThermoFisher). RT-qPCR was carried out on a Quantstudio 6 flex system (ThermoFisher) using PerfeCTa SYBR Green FastMix (Quantabio) and the appropriate primers for each target gene (Table S1).

### Quantifying the area of potato consumed by larvae in 24 hours

To assess how temperature affects consumption, larvae were trapped inside clip-cages on four-week-old Kennebec potato plants and incubated at either 18, 25 or 30°C. For the between-cultivar consumption analysis, larvae were trapped on four-week-old Kennebec, Chieftain and Purple Viking potato plants and incubated at 25°C. All potato plants were grown in six-inch pots as described above. In the field, CPB larvae generally hatch on the abaxial surface of leaves, and sprayed dsRNA will contact the adaxial surface. To mimic field conditions, leaf treatments were applied to the adaxial surface of leaves, and first instar larvae were trapped on the abaxial surface. 15 leaves on each plant had 10 µl dsRNA (1:1, 900 ng/µl dsactin: Triton-X (53nM)) and 10 µl water (1:1, H2O: Triton-X (53 nM)) pipetted onto opposite sides of the mid-vein, giving a total of 30 feeding sites for each plant. Larvae were then trapped at the feeding sites in custom-made clip cages. Plants were moved to an incubator at the appropriate temperature (16L:8D) with the roots positioned in a reservoir of water for 24 hours. The clip cages were removed, leaving a 12 mm Ø impression of the feeding site on the leaf. The unconsumed leaf tissue remaining at each feeding site was then cut from the plant with a 12 mm Ø hole punch. The leaf tissue was then imaged at 2400 dpi with a perfection V800 scanner (Epson) and the area was quantified using FIJI v1.54j.

### Quantification of the metabolites contained in leaf tissue

12mm Ø leaf discs were cut from 4-week-old Kennebec, Chieftain and Purple Viking potato plants. 10 mature leaves were selected from two replicate plants per cultivar, and a single disc was cut from each. The discs were homogenized, and total protein, carbohydrate and lipids were quantified according to Foray et al. (2012).

**1) Homogenisation and aliquoting** – Discs were homogenised in 360 µl of lysis buffer (100 mM KH_2_PO_4_, 1 mM DDT, 1 mM EDTA) using four 2.3 mm zirconia/silica beads (Biospec) and a FastPrep-24 bead beater (MP). The homogenised samples were centrifuged at 180 g and 4°C for 15 mins, and 180 µl of the supernatant was aliquoted for the Bradford protein assay (see below). The remaining homogenate had 20 µl of 20% sodium sulphate added to it and was then vortexed. 1500 µl of 1:2 Chloroform: Methanol was added to the homogenate, which was vortexed again before being centrifuged at 180 g and 4°C for 15 mins. 300 µl of the supernatant was aliquoted into a glass tube for lipid quantification (see below) and incubated at 90°C until the solvent had evaporated. For downstream analysis of soluble carbohydrates (see below), 150 µl of the supernatant was aliquoted into a glass tube and incubated at 70°C for one hour until solvents had evaporated. The remaining supernatant was discarded, and the pellet was washed with 400 µl of 80% ethanol, vortexed and centrifuged at 16000 g and 4°C for 5 mins. The wash step was repeated, supernatants were removed by pipette and the pellet was retained for starch analysis (see below).
**2) Bradford protein assay** – To quantify proteins, we used the Bradford assay with an eight-point albumin (ThermoFisher) standard curve ranging from 0-2000 µg/ml. Supernatants that were aliquoted for Bradford analysis during the homogenisation step were diluted four-fold, and duplicate 5 µl aliquots of each sample were pipetted into a 96-well microplate (Greiner). Duplicate 5 µl aliquots of the albumin standard curve (range = 0-10 µg total protein) were then pipetted into the same microplate. 250 µl of Coomassie reagent (ThermoFisher) was added to all samples and standards, and the contents of the plate were mixed by shaking for 30 secs, then incubated at room temperature for 10 mins. Absorbance at 595 nm was measured using a CLARIOstar plate reader (BMG) and the mass of protein contained in leaves was calculated by linear regression, using the standard curve.
**3) Lipid analysis** – To quantify lipids, we used a vanillin assay with an eight-point standard curve of vegetable oil (0-300 µg). Vegetable oil was dissolved in chloroform and incubated at 90°C until the solvent had evaporated. All homogenised leaf disc samples and the standards then had 100 µl of sulphuric acid (VWR) added, before being vortexed and incubated at 90°C for 10 mins. 1 ml of vanillin reagent (1.2 mg/ml of vanillin (VWR) in 68% phosphoric acid (VWR)) was added to the tubes, before they were vortexed and incubated at room temperature for 20 mins. Duplicate 100 µl replicates were pipetted from each tube into a 96-well microplate (Greiner). Absorbance was measured at 525 nm using a CLARIOstar plate reader (BMG) and the mass of lipid contained in leaves was calculated using the standard curve, by linear regression.
**4) Soluble carbohydrate and starch analyses** – To quantify soluble carbohydrates and starch, we used the anthrone assay with an eight-point standard curve of D-glucose (0-2000 µg/ml) (ThermoFisher). 100 µl of water was added to the soluble carbohydrate and starch samples that were kept aside during the homogenisation. The samples were vortexed and incubated at 70°C for 5 mins to dissolve the sugars. 400 µl of anthrone reagent (1.4 mg/ml anthrone (AlfaAesar) in 72% sulphuric acid (VWR)) was added to all tubes containing samples and standards, giving a total volume of 500 µl in each tube. The tubes were vortexed and incubated at 100°C for 17 mins, then vortexed again. Duplicate 125 µl replicates were pipetted from each tube into a 96-well microplate (Greiner). Absorbance was measured at 625 nm using a CLARIOstar plate reader (BMG) and the mass of soluble carbohydrate and starch contained in leaves were calculated using the standard curve, by linear regression.

### Statistics

Data were consolidated and transformed using the Tidyverse (Wickham, 2017) in R (Team, 2016). Data were then modelled and analysed using the survival (Therneau, 2020), lme4 (Bates et al., 2015), car (Fox & Weisburg, 2019) and emmeans (Lenth, 2024) packages in R (Team, 2016).

## Acknowledgements

We thank Jack Graba for assistance with plant and insect husbandry.

## Data availability statement

The data that support the findings of this study are openly available in github https://github.com/miked198044/RNAi-in-varying-environments

